# TEX-FBA: A constraint-based method for integrating gene expression, thermodynamics, and metabolomics data into genome-scale metabolic models

**DOI:** 10.1101/536235

**Authors:** Vikash Pandey, Daniel Hernandez Gardiol, Anush Chiappino-Pepe, Vassily Hatzimanikatis

## Abstract

A large number of genome-scale models of cellular metabolism are available for various organisms. These models include all known metabolic reactions based on the genome annotation. However, the reactions that are active are dependent on the cellular metabolic function or environmental condition. Constraint-based methods that integrate condition-specific transcriptomics data into models have been used extensively to investigate condition-specific metabolism. Here, we present a method (TEX-FBA) for modeling condition-specific metabolism that combines transcriptomics and reaction thermodynamics data to generate a thermodynamically-feasible condition-specific metabolic model. TEX-FBA is an extension of thermodynamic-based flux balance analysis (TFA), which allows the simultaneous integration of different stages of experimental data (e.g., absolute gene expression, metabolite concentrations, thermodynamic data, and fluxomics) and the identification of alternative metabolic states that maximize consistency between gene expression levels and condition-specific reaction fluxes. We applied TEX-FBA to a genome-scale metabolic model of *Escherichia coli* by integrating available condition-specific experimental data and found a marked reduction in the flux solution space. Our analysis revealed a marked correlation between actual gene expression profile and experimental flux measurements compared to the one obtained from a randomly generated gene expression profile. We identified additional essential reactions from the membrane lipid and folate metabolism when we integrated transcriptomics data of the given condition on the top of metabolomics and thermodynamics data. These results show TEX-FBA is a promising new approach to study condition-specific metabolism when different types of experimental data are available.

**Author summary:** Cells utilize nutrients via biochemical reactions that are controlled by enzymes and synthesize required compounds for their survival and growth. Genome-scale models of metabolism representing these complex reaction networks have been reconstructed for a wide variety of organisms ranging from bacteria to human cells. These models comprise all possible biochemical reactions in a cell, but cells choose only a subset of reactions for their immediate needs and functions. Usually, these models allow for a large flux solution space and one can integrate experimental data in order to reduce it and potentially predict the physiology for a specific condition. We developed a method for integrating different types of omics data, such as fluxomics, transcriptomics, metabolomics into genome-scale metabolic models that reduces the flux solution space. Using gene expression data, the algorithm maximizes the consistency between the predicted and experimental flux for the reactions and predicts biologically relevant flux ranges for the remaining reactions in the network. This method is useful for determining fluxes of metabolic reactions with reduced uncertainty and suitable for performing context- and condition-specific analysis in metabolic models using different types of experimental data.

## Introduction

The study of cellular metabolism has important applications in medicine, biology, and biotechnology. Genome-scale metabolic models (GEMs) are useful scaffolds to integrate “omics” data and analyze the metabolic capabilities of organisms and their metabolic pathways giving rise to different physiologies [1]. In particular, GEMs have been successfully used for engineering strains for optimizing the production of chemicals [2-4] and identifying drug targets for metabolic diseases, such as diabetes [5], cancer [6-8] and non-alcoholic fatty liver disease [9, 10]. Increased availability of organisms’ gene sequencing has led to an accumulation of available GEM reconstructions for a variety of bacteria and eukaryotic organisms [11-20].

GEMs are amenable to the integration of different types of “omics” data (e.g., fluxomics, metabolomics, and transcriptomics) to simulate condition- and organism-specific metabolic phenotypes, which allows to contextualize the observed cellular metabolism according to its environmental conditions and intrinsic metabolic capabilities. Flux Balance Analysis (FBA) is the most commonly used method that can integrate uptake and secretion rates, as well as intracellular fluxes, if known, to constrain the cellular physiology and simulate condition-dependent metabolic phenotypes [21-23].

Other methods have extended FBA to allow integration of intracellular metabolite concentrations and predict thermodynamically feasible cellular metabolic states [24-27]. GEMs are often reconstructed with pre-assigned directionalities imposed to the reactions from extensive literature curation on biochemical assays that may not hold for all cellular conditions. However, for catalytically reversible reactions, the direction of the net flux is determined by thermodynamic properties that also depend on the intracellular concentration of products and reactants, which themselves are condition-dependent. Advances in group contribution method for the estimation of thermodynamic properties of metabolites [28] and the development of thermodynamics-based flux balance analysis (TFA) [24-26] allow for the reaction thermodynamic properties along with metabolomics data to be integrated into GEMs. Accounting for thermodynamics and metabolomics ensures that the metabolic fluxes operate within physiologically realistic confines.

The technological advances in mRNA sequencing techniques increased the access to transcriptomics data. As a result, several FBA-based algorithms have been developed to predict metabolic fluxes based on absolute [29-33] or relative [34-37] gene expression data, which can be measured for different tissues and conditions. Among the methods that use absolute transcriptomics data, the Lee et al. [31] and iMAT [32, 33] methods do not require assumptions on cellular objective functions, which is an advantage for the study of organisms whose cellular metabolic function is not evident, such as for multicellular organisms. However, while the formulation of the Lee et al. method estimates the fluxes of a cellular metabolic state by minimizing their differences with respect to the gene expression levels, iMAT allows for the investigation of alternative flux distributions that are consistent with the transcriptomics data. Upon integration of the gene expression data and its association with the respective enzymatic reactions in the GEM, iMAT maximizes and minimizes the number of highly- and lowly-expressed reactions, respectively, returning a GEM whose fluxes are constrained to a metabolic state that is most consistent with the data. To our knowledge, no previous method has simultaneously integrated expression data, metabolic concentrations, and thermodynamic constraints into GEMs. Here, we present TEX-FBA, an algorithm developed by extending the iMAT formulation to allow integration of both thermodynamics and metabolomics constraints simultaneously with transcriptomics with the purpose of generating a context-specific thermodynamically-feasible model without imposing *a priori* directionalities on the fluxes. TEX-FBA assumes a correlation between absolute gene levels and reaction fluxes, and it enforces a high and a low flux for the reactions associated with highly- and lowly-expressed genes, respectively.

In this study, we applied TEX-FBA to generate a condition-specific model of *Escherichia coli* (*E. coli*) that integrates for the first time different types of omics data [38], such as transcriptomics, fluxomics, and metabolomics as well as thermodynamic data into an *E. coli* GEM [39]. We used this model to study reaction flux ranges that are resulted by the model integrated with gene expression data. When we compared the model-resulted flux ranges with experimental flux measurements we found that actual expression data are markedly consistent with flux measurements as compared to a random gene expression profile. TEX-FBA identifies alternative states of optimal consistent constraints from gene expression data, which makes TEX-FBA different than iMAT. As expected, we found a decrease in flux solution space and uncertainty after integrating metabolomics and thermodynamics data as compared to integrating only fluxomics and transcriptomics data. We identified essential genes and reactions that are required for the cell’s growth. Only the simultaneous integration of metabolomics and thermodynamics data resulted in the identification of essential reactions from central carbon metabolism. By further integrating gene expression data we identified additional essential reactions from folate and lipid membrane metabolism. TEX-FBA method that can be widely applicable to study condition-specific metabolism for other organisms based on different types of omics data.

## Results and Discussion

### A sensitivity analysis of the reaction flux ranges based on the TEX-FBA parameters and its comparison with C13 fluxomics data

We developed the TEX-FBA method for integrating thermodynamics, transcriptomics, metabolomics, and fluxomics into GEMs in this study. TEX-FBA has four parameters (see Materials and Methods): two (*l, h*) are used for classifying genes into lowly- and highly-expressed gene categories, respectively, and the reaming two parameters (*p*_*l*_, *p*_*h*_) are flux constraints used to enforce flux through reactions corresponding to lowly- and highly-expressed genes, respectively. Values of these parameters are user defined. Then, we applied TEX-FBA to the *E. coli* GEM iJO1366 [39] with condition-specific gene expression data of glucose-limited *chemostat E. coli* cultures at a dilution rate of 0.2 h^−1^ [38] and thermodynamics constraints for given parameter values and estimated flux ranges.

The parameter *l* was varied between 5 to 20 percentile by setting the values: 5, 10, 15, and 20, and similarly we also varied the parameter *h* from 80 to 95 percentile with the values: 80, 85, 90, 95. The parameter *p*_*h*_ was set with the values: 0.6, 0.7, 0.8, and 0.9. A value of 0.6 for *p*_*h*_ means that the flux is allowed to vary above 60% of the flux variability range (see Materials and Methods). We set the parameter *p*_*l*_ with the values: 0.1, 0.2, 0.3, and 0.4. A value of 0.1 for *p*_*l*_ means that the flux is allowed to vary below 10% of the flux variability range. In total, 256 (4^4^) parameter combinations were used for a sensitivity analysis of the TEX-FBA method. For each parameter combination, we obtained a maximum consistency score (MCS; see Material and Methods) that measures the maximum number of feasible constraints from expression data. TEX-FBA allows for the enumeration of alternative flux profiles (Table S1) for a given MCS due to the possibility of combining various sets of fluxes constrained from expression data that lead to the same MCS (see Materials and Methods). For example, for a parameter set (*l*=10, *h*=95, *p*_*l*_=0.1, *p*_*h*_=0.6), we obtained MCS of 70 with 4 alternatives. In total for MCS, we obtained 412 alternatives with 256 parameter combinations (Table S1). We generated 412 alternative models (see Materials and Methods, generation of alternative models) by activating expression constraints, and performed Flux Variability Analysis (FVA) [40] for each alternative model.

The flux ranges of the 34 reactions for which we have C13 measurements [38] were compared to the flux ranges obtained with FVA using the Flux Range Comparisons metric (FRC; see Materials and Methods). For all measured 34 C13-based flux ranges, we first calculated FRCs for each reaction and then calculated an average FRC (Table S2). The average FRC scores were ranked for each alternative model (Table S2). The lowest average FRC score was 286.6 for the parameter values *l*=15, *h*=85, *p*_*l*_=0.3, and *p*_*h*_=0.7 (Table S2). The parameter set rendering the lowest average FRC was thus the best fit to the condition-specific expression and C13-based flux data. To test for the specificity of the obtained best-fit parameters, we generated 100 random gene expression profiles, and each random profile was integrated into the iJO1366 using TEX-FBA method and best-fit parameter. Similar to the TEX-FBA with actual gene expression, we also performed maximum consistency and alternative enumeration analysis of the 100 random gene expression profiles and obtained 220 alternatives. The lowest FRC score obtained when using the expression profile measurements was significantly lower (p-value=1.8 e-48) than the obtained from the random expression profiles (average FRCs=3.74). Therefore, we can safely conclude that the actual gene expression profile outperformed the random profile in predicting a flux range close to the C13 experimental measurements.

We next searched for patterns in parameters that agreed with C13-based flux measurements. We generated a random set of average FRCs between zero and a large number, where zero indicates a predicted flux range agrees with C13-based flux measurement and a large number shows disagreement between prediction and measurement. We compared the random profile of average FRCs with the average FRCs of the 412 alternative models, which integrated actual gene expression data. We identified 94% out of 412 alternative models that have significantly low FRC score (p-value < 1e-05) compared to an average of random FRCs (Table S2), which means these models were in good agreement with C13 flux measurements. Interestingly, only a few alternative models (6%) disagreed with C13 flux measurements and performed like random. We next identified ranges of parameter values that frequently occurred in the population of models that showed high average FRCs and might hence be responsible for the disagreement with fluxomics data. For example, in the models that disagreed with fluxomics data the parameter value *p*_*h*_=0.9 was found to occur frequently. The higher value of *p*_*h*_ (*p*_*h*_=0.9) indicates that enforcing a very high flux (90% of a flux range) through the highly expressed reactions resulted in a poorer agreement with experimental fluxomics data. Based on these results, we hypothesize that enforcing very high flux even for a small number of highly-expressed reactions might reroute fluxes through metabolic networks, unbalance enzyme usage, and result in poor agreement with omics data, which resonates with previous observations [41]. Interestingly, enforcing a moderate flux (*p*_*h*_=60% to 70% of a flux range) through highly expressed reactions yields flux predictions that agree better with fluxomic measurements. This result suggests that a context-specific metabolic network maintains moderate flux levels for it highly-expressed reactions, which might allow a faster flux redistribution and survival upon perturbation.

### Generation of four models that integrate different combinations of datasets for the study of context-specific metabolism

We used the best-fit parameter values: values *l*=15, *h*=85, *p*_*l*_=0.3, *p*_*h*_=0.7 for integrating expression data. Starting from the *E. coli* GEM iJO1366 [39], we generated four models corresponding to different stages of data integration: a FBA model, a TFA model, a XFBA model, and a XTFA model (Figure 1). The data used consisted of expression data, intracellular metabolite concentrations, and flux data collected from experiments performed in glucose-limited *chemostat E. coli* cultures at a dilution rate of 0.2 h^−1^ [38]. The *FBA model* was generated by integrating only the flux data, whereas the *TFA model* was further constrained with thermodynamic constraints and intracellular concentrations using the TFA method [25]. We then applied the TEX-FBA formulation to integrate expression data using the best-fit parameter and generated the XFBA and XTFA models (see Materials and Methods). The *XFBA model* was generated with transcriptomics data integrated simultaneously with the flux data but without applying thermodynamic constraints and intracellular metabolite concentrations. The *XTFA model* was generated by integrating fluxomics, thermodynamic constraints applied to the reactions, intracellular metabolite concentrations, and transcriptomics data simultaneously.

**Figure 1.**
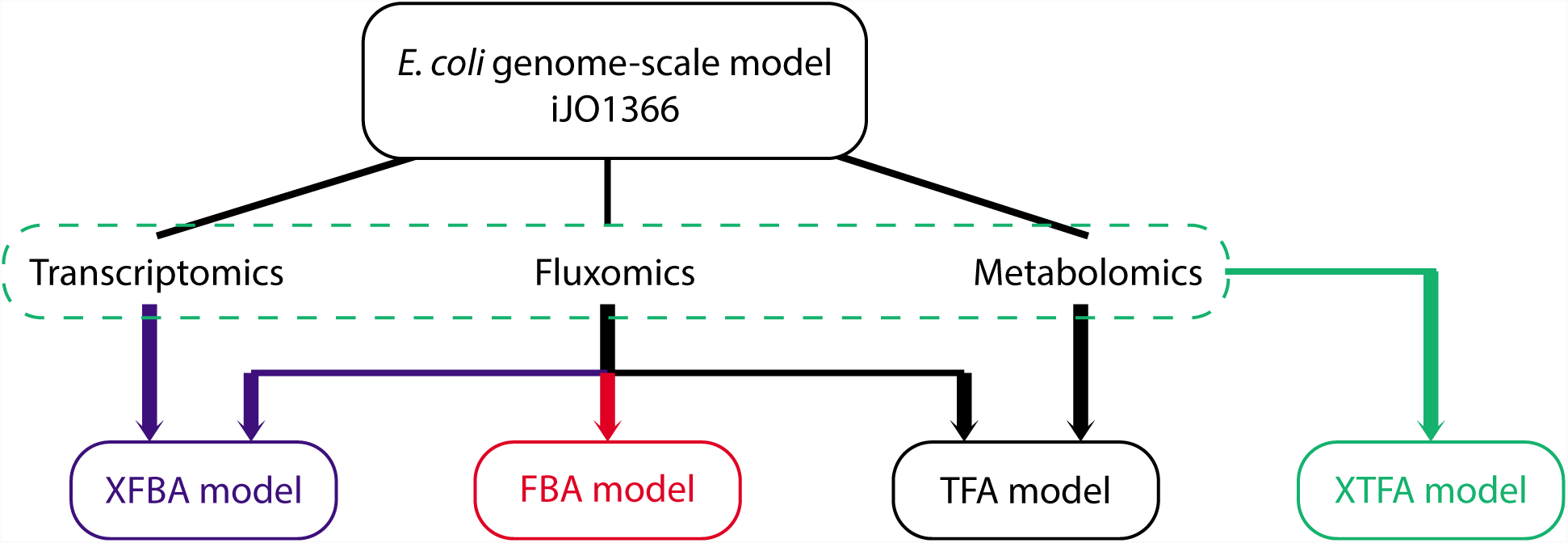
Workflow of model generation in this study. Boxes with a solid line indicate models at different stages of data integration. At the top, we find the original *E. coli* genome-scale model iJO1366 [39]. At the bottom, we present four models generated in this study: the FBA model (red), XFBA model (blue), TFA model (black), and XTFA model (green). Arrows indicate the flow of input data (transcriptomics, fluxomics, and metabolomics from [38]) to generate each model.

### Analysis of highly- and lowly-expressed reactions in alternative profiles at maximum consistency in XFBA and XTFA models

We used the XFBA and XTFA models generated to identify highly- and lowly-expressed genes and their associated highly- and lowly-expressed reactions. Reactions with assigned gene expression data are called Expression Constraint Reactions (ECRs). We identified such 259 ERCs, which represent the Maximum Theoretical Consistency Score (MTCS; see Materials and Methods). After a TEX-FBA without (XFBA model) and with (XTFA model) thermodynamic constraints we found a value of 81 and 75 for the MCS, respectively. The MCSs values of 81 and 75 indicate that out of 259 ECRs we can obtain maximally 81 reactions (31%) and 75 reactions (29%) consistent with gene expression for the XFBA and XTFA models, respectively. For any consistency score, there can be alternative consistency states, defined by the combination of active expression binary variables (see Materials and Methods). For the given MCSs of the XFBA and XTFA models, we obtained three and two alternative consistency states, respectively.

We next analyzed the frequency with which a reaction is consistent with gene expression over all alternatives, which is an indication of the certainty of that prediction. There were 71 reactions (27% of the 259 ECRs) that were consistent with gene expression across all alternatives of the XFBA and XTFA models (Table S3). All 71 reactions were lowly expressed. We performed a subsystem enrichment analysis on the 71 reactions (see Materials and Methods) to identify the subsystems that were consistently lowly expressed in the context of the study. The metabolic subsystems that were significantly enriched (p-value < 0.05) with the 71 lowly-expressed reactions were the metabolism of alternate carbon, folate, and glycoxylate, and tricarboxylic acid cycle (TCA) (Table S3). TCA was also enriched by highly expressed reactions, meaning that different control of fluxes appears through the TCA (Table S3). Interestingly, previous studies have shown that the control of flux through TCA in *E. coli* allows to sustain aerobic growth [42].

Conversely, 174 reactions (67% of the 259 ECRs) were inconsistent with gene expression across all alternatives (Table S3). All were highly-expressed reactions that significantly enriched the cell envelope biosynthesis pathway (p-value<0.05; Table S3). For the 174 reactions, the predicted amount of flux was not as high as expected (*p*_*h*_) given their high transcript expression. The lack of correlation between gene and flux levels suggests there might exist post-transcriptional regulation, as suggested before [32], or the enzymatic activities of the enzymes might be higher than the average.

The remaining 14 reactions (5% of the 259 ECRs) showed different consistency with gene expression across alternatives and between the XFBA and XTFA models (Figure 2). We identified some reactions in the membrane lipid metabolism such as HACD7 and ECOAH7 that were uniquely active in the XFBA model (Figure 2). Such XFBA-specific reactions should not be consistent with gene expression based on thermodynamic constraints. This result suggests one should account for thermodynamics to avoid erroneous conclusions on the context-specific physiology. Some reactions of the alternate carbon metabolism such as PPCSCT (Figure 2) were uniquely active in the XTFA model. Such reactions could become consistent with gene expression thanks to thermodynamic constraints. The reaction EDTXS3 in lipopolysaccharides biosynthesis was switched between the alternatives from the XFBA and XTFA models (Figure 2), indicating that a low flux through these reactions is valid for both models with or without reaction thermodynamics, but this result is alternative-dependent. This observation demonstrates the importance of analyzing alternative optimal solutions for drawing reasonable conclusions on the context-specific metabolic function, as suggested before [43].

**Figure 2.**
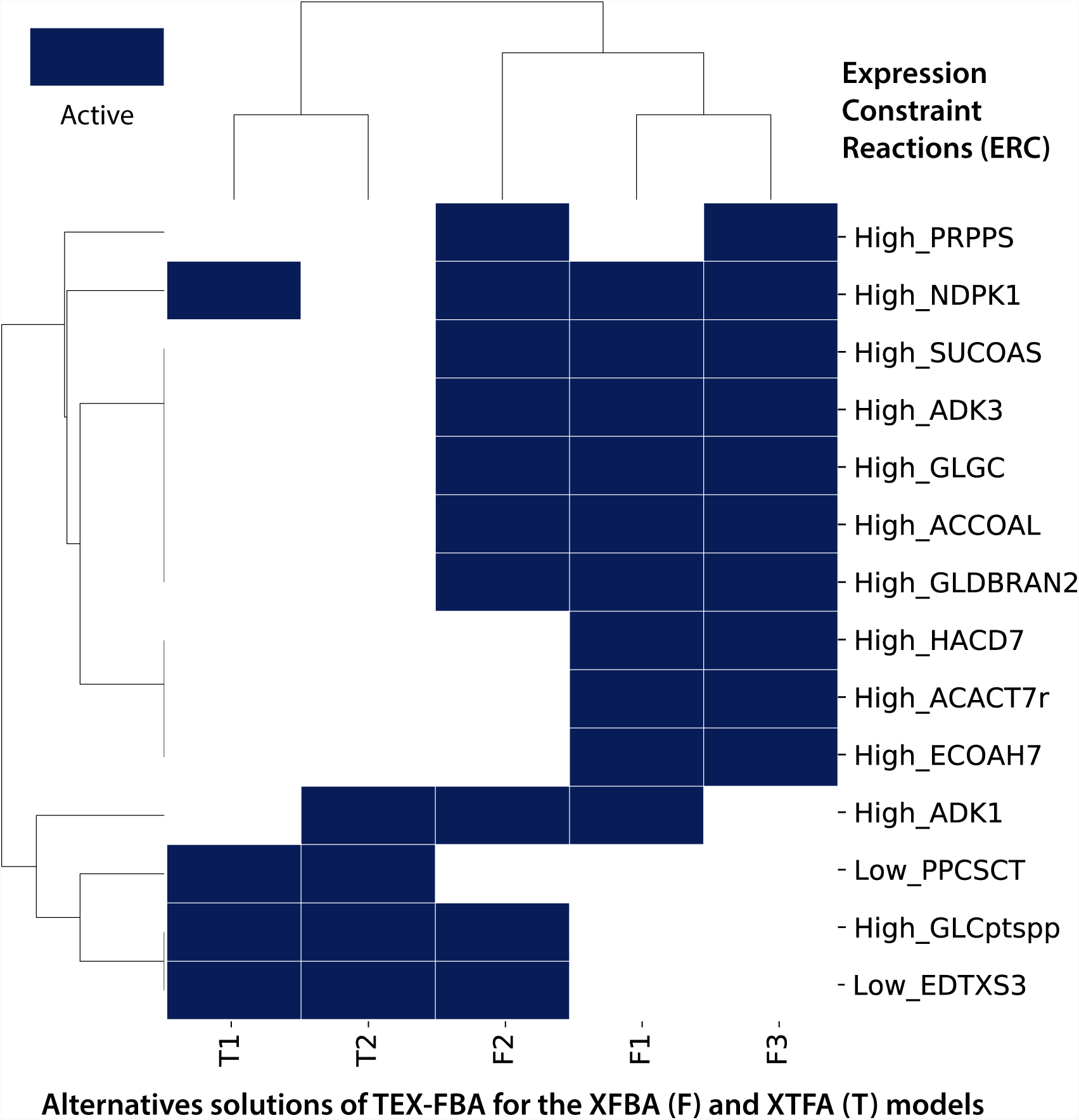
Clustering of alternative TEX-FBA solutions (OP 1) at Maximum Consistency Score (MCS). We show two and three alternative solutions of the XTFA (columns with a capital T) and XFBA models (columns with a capital F), respectively. Rows indicate Expression Constraint Reactions (ECRs). The ECRs are labeled with “high” or “low” based on the reaction level assigned with the TEX-FBA methodology (see Materials and Methods). The ECRs are presented with the reaction IDs as provided in the *E. coli* GEM iJO1366 [39]. Dark blue indicates that an ECR is active in the alternative solution (*y*_*i*_ = 1 in the OP 1), which means it is consistent with gene expression.

### Analysis of reduction of uncertainty on models with different datasets integrated indicate the importance of thermodynamic constraints

The integration of omics data into genome-scale models allows the study of context-specific metabolism through a reduction of uncertainty in the estimation of fluxes. We performed a flux variability analysis (FVA) [40] to determine the feasible reaction flux ranges (here referred to as flexibility) of the four models (FBA, XFBA, TFA, and XTFA) generated in this study. The FVA results of the XFBA, TFA, and XTFA models compared to the FBA model served to quantify a reduction of flexibility by integrating different types of data. We computed a relative flexibility metric per reaction and an average relative flexibility (ARF) per metabolic subsystem and model (see Materials and Methods). We also identified the number of blocked reactions, i.e., reactions that cannot carry flux, and bidirectional reactions, i.e., reactions that can carry flux in both forward and backward directions in each model.

Out of the three models studied, XTFA showed the lowest ARF compared to the FBA model (Figure 3A). This result suggested that the difference in flux ranges between the FBA and XTFA model was the highest one, which was expected because the XTFA model integrated all datasets [39] (Figure 1) and thermodynamic constraints that eliminated thermodynamically infeasible flux solutions. The XTFA model was hence more constrained than the FBA, XFBA, and TFA models and allowed the analysis of context-specific metabolism and fluxes with the highest reduction of uncertainty. The study of reaction directionality supported this result. The XTFA model presented the lowest number of bidirectional reactions (Table 1). There were 56 bidirectional reactions in the FBA model that became unidirectional in the XTFA model (Table S4).

**Table 1.** Number of blocked and bidirectional reactions in the FBA, TFA, XFBA, and XTFA models.

| Models | Nb of blocked reactions | Nb of bidirectional reactions |
| --- | --- | --- |
| FBA | 915 | 124 |
| TFA | 1,273 | 73 |
| XFBA | 1,537 | 95 |
| XTFA | 1,267 | 69 |

**Figure 3.**
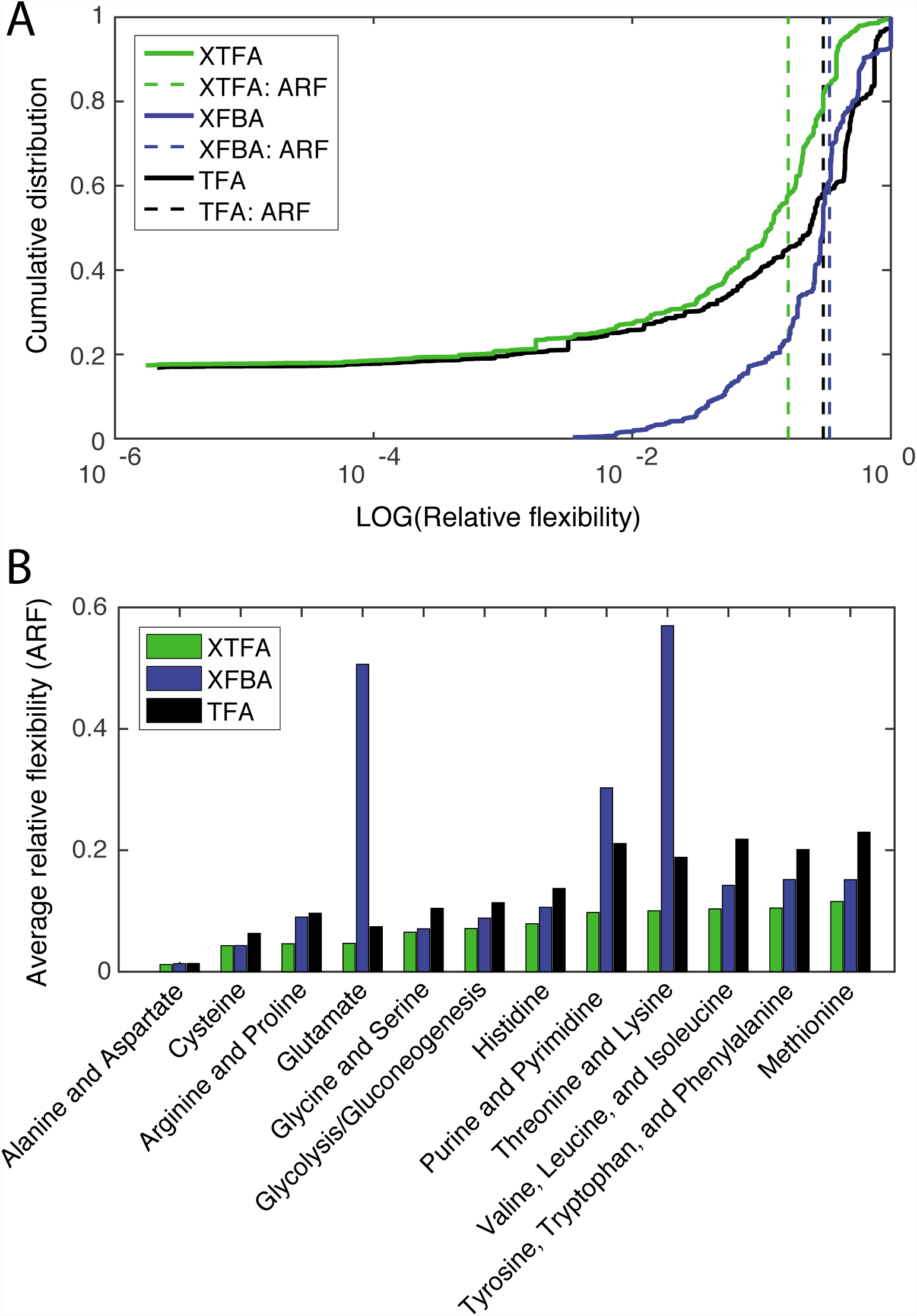
Results of the analysis of relative flexibility on the XFBA (blue), TFA (black), and XTFA (green) models compared to the FBA model. (A) We show the cumulative values of the relative flexibility metric per reaction in the three models with solid lines. We also depict the average relative flexibility of the XFBA, TFA, and XTFA models with a vertical dashed line. (B) We present the relative flexibility per metabolic subsystem in the three models. We show the top twelve metabolic subsystems that show the lowest relative flexibility for the XTFA model.

The ARF of the XFBA and TFA models was close to each other (Figure 3A). This result indicated that integration of the available gene expression dataset [39] (XFBA model) or thermodynamic constraints (TFA model) into the *E. coli* metabolic model reduced the flux solution space and uncertainty to a similar extent in the analysis of condition-specific physiologies. Interestingly, we identified 24 reactions that were initially bidirectional in the FBA model and became unidirectional in both XFBA and TFA models (TableS4). This result further suggests the consistency in constraining the iJO1366 with both gene expression and thermodynamic constraints. Availability of a bigger set of metabolomics data or concentration data for some key metabolites (as suggested before [12, 44]) might have reduced the relative flexibility of the TFA model and increased the difference between the XFBA and TFA relative flexibility compared to the FBA model.

Analysis of ARF per metabolic subsystem highlighted a reduction of flexibility in the amino acid metabolism, glycolysis, and purine/pyrimidine biosynthesis of the TFA model (Figure 3B). All metabolic subsystems of the XTFA model showed the highest reduction of ARF relative to the FBA model, which suggested the reduction of ARF observed in the XTFA model was a consequence of a similar decrease of uncertainty throughout the metabolic network. Despite showing similar ARF per model, the ARF per metabolic subsystem of the XFBA and TFA models was significantly different with a particularly heterogeneous reduction of uncertainty for the FBA model (Figure 3B).

### Analysis of condition-specific essentiality on models with different datasets integrated demonstrate the relevant impact of thermodynamic constraints

The metabolic functions that are essential for growth vary with the environmental and genetic background of the cell [45]. We performed gene and reaction essentiality analysis on the four models with different stages of data integration to study condition-specific essentiality (see Materials and Methods). We identified 385 essential reactions and 282 essential genes in the FBA model, which were indispensable also in the XFBA, TFA, and XTFA models (Figure 4 and Tables S5-S6) since these models integrated additional data and constraints (Figure 1). The TFA model showed 14 more essential reactions and five more essential genes than the FBA model, as a product of thermodynamic constraints. Gene expression on the top of thermodynamic constraints resulted in two new essential reactions and no new essential genes in the XTFA model compared to the TFA model. We identified eleven reactions and twelve genes that were essential uniquely in the XFBA model and are hence the result of gene expression constraints that are not thermodynamically allowed (Figure 4 and Tables S5-S6).

**Figure 4.**
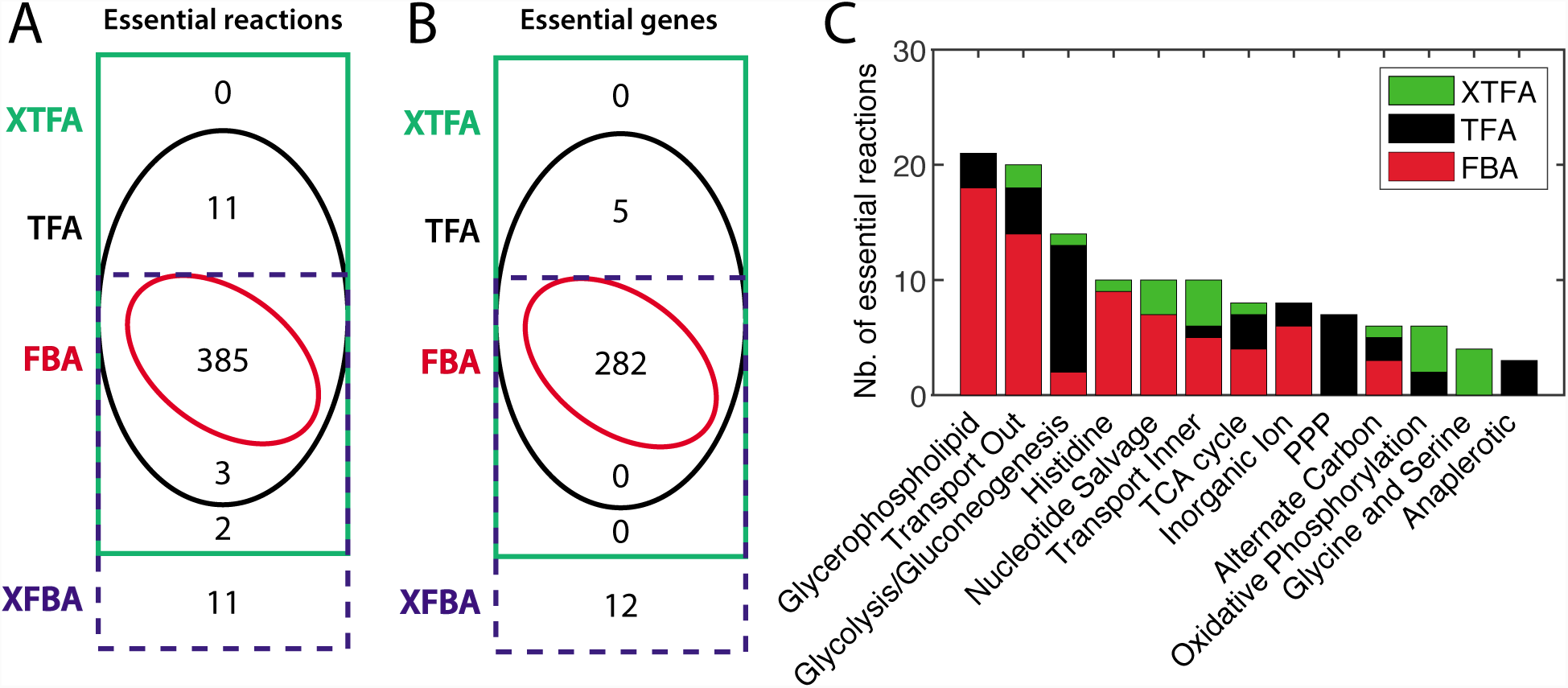
Results of gene and reaction essentiality analysis on the FBA (red), XFBA (blue), TFA (black), and XTFA (green) models. (A) Overlap in the prediction of essential reactions between the models. (B) Overlap in the prediction of essential genes between the models. (C) Distribution of essential reactions per metabolic subsystem in the FBA, TFA, and XTFA models. We show the top thirteen subsystems with the highest number of essential reactions in the XTFA model.

We further identified the metabolic subsystems associated with context-specific essentiality. Interestingly, thermodynamic constraints were crucial for identifying essential reactions in the central carbon metabolism: glycolysis, TCA, and pentose phosphate pathway (PPP) (Figure 4B; black). This result suggests that accounting for thermodynamic and metabolite concentration data eliminates redundancy through thermodynamic infeasibility in the central carbon metabolism. The integration of gene expression into the TFA model resulted in essential transporters and reactions in the lipid, folate, and amino acid metabolism (Figure 4B; green). To our knowledge, there are no experimental data on gene essentiality at the conditions of study for comparison. However, we hypothesize these genes are essential for growth in the studied conditions given the high consistency of the XTFA model with fluxomics data (as shown previously in this result section).

## Conclusions

The study of context-specific metabolism requires methods that allow the integration of different kinds of datasets into GEMs. The lack of methodologies that enable accounting for thermodynamics, metabolomics, and gene expression data simultaneously motivated us to develop TEX-FBA. As all methods for integrating gene expression data into GEMs, TEX-FBA assumes that mRNA transcript levels are strongly correlated with levels of protein activity and reaction fluxes. However, post-transcriptional regulation and post-translational modification can affect the final protein activity, and several studies in yeast have shown that direct correlation between transcripts and protein levels [46, 47] or metabolic fluxes [48] can be rather weak. TEX-FBA allows for the presence of low correlation between mRNA and flux levels by trying to maximize and not imposing such correlation. TEX-FBA defines a consistency score that allows to define a desired level of agreement between gene expression and fluxes. Moreover, TEX-FBA identifies alternative states of consistency between gene expression and metabolic fluxes, which makes this method unique to others. The analysis of alternative optima is indeed required to reduce the uncertainty of context-specific metabolic model predictions, as described before [43] and shown in this study.

The conclusions that might arise from context-specific metabolic modeling for a physiology will depend on the parameter values of the method used. We have suggested a sensitivity analysis procedure to evaluate the impact of the parameters on the flux values and identify best-fit parameters for TEX-FBA.

We integrated fluxomics, metabolomics, and gene expression data of *E. coli* [38] into a GEM of this organism and generated four models with various stages of data integrated (Figure 1). We demonstrated that the simultaneous integration of all datasets markedly reduces the uncertainty in the estimation of fluxes and reaction directionalities (Table 1), which is a valuable input for further studies such as kinetic analysis [49]. We also identified a strikingly higher number of essential genes and reactions in the *E. coli* GEM when all omics data were taken into account. Interestingly, there was a significant number of condition-specific essential reactions for growth in the central carbon metabolism that only thermodynamic constraints were able to identify. We, therefore, believe the TEX-FBA method is widely applicable and compelling for studying context-, condition-, and life-stage-specific metabolism for any given organism or disease.

## Methods

### Organism, genome-scale model, and physiological data

We used the genome-scale model of *E. coli* K-12 iJO1366 [39] to generate context-specific GEMs using different data types. We integrated three data sets into iJO1366: (i) absolute gene expression data from microarrays, (ii) intracellular metabolite concentrations, and fluxomics data, i.e., estimated uptake and secretion rates. All the data was collected from experiments performed in glucose-limited chemostat cultures at a dilution rate of 0.2 h^−1^ [38]. See the software section for more details on the availability of the model and data used.

### The TEX-FBA formulation and Maximum Consistency Score (MCS)

The lack of a methodology that allows integrating fluxomics, thermodynamic constraints, metabolomics, and transcriptomics simultaneously into a GEM motivated us to develop TEX-FBA. The TEX-FBA formulation is based on the iMAT methodology [32, 33] and extends it with the integration of thermodynamic constraints [25, 26, 50] and a new definition of flux ranges consistent with gene expression. The implementation of TEX-FBA follows a sequence of eight steps:

1. We compute the reaction flux levels based on the associated gene expression levels and the gene-protein-reaction (GPR) rules of the GEM. We make the following assumptions to treat the reactions catalyzed by (i) an enzyme complex, i.e., genes associated with an AND rule like geneA AND geneB, or (ii) by isoenzymes, i.e., genes associated with an OR rule like geneA OR geneB. If there are lowly and highly expressed genes together in an (i) AND or (ii) OR rule, we compute the associated reaction flux level by (i) taking the minimum gene level or (ii) adding the gene levels, respectively. We follow the FALCON methodology implemented by Baker and colleagues [51] to account for protein subunit levels in enzyme complexes and estimated reaction flux levels.
2. We then classify the reactions in three groups, i.e., lowly-, moderately-, and highly-expressed, based on the computed flux level. Two parameters *l* and *h* divide the distribution of reaction flux levels into the three states. While in the original iMAT formulation *l* and *h* are user-defined, in the TEX-FBA formulation they represent the bottom and top percentile of the computed reaction expression values, respectively. We classify a reaction as lowly- (*r*_*L*_), moderately- (*r*_*M*_) or highly- (*r*_*H*_) expressed if its calculated expression level is below *l*, between *l* and *h*, or above *h*, respectively.
3. As done within the TFA framework, we split the reaction fluxes in forward and backward to work with positive flux values. We also associate the forward and backward fluxes to integer variables and assure they are not active simultaneously.
4. We perform a flux variability analysis (FVA) [40] to determine the reaction flux ranges *FR* = [*v*_*min*_, *v*_*max*_], where *v*_*min*_ and *v*_*max*_ is the minimum and maximum flux value in the range.
5. We define a maximum and minimum flux requirement for the lowly and highly expressed reactions, respectively, by setting flux constraints within the flux range. For the highly-expressed reactions, we define *p*_*h*_ to increase the *v*_*min*_ and constrain the flux range to *FR* = [*p*_*h*_ · (*v*_*max*_ − *v*_min_), *v*_*max*_]. For the lowly-expressed reactions, we set *p*_*l*_ to reduce the *v*_*max*_ and restrict the flux range to *FR* = [*v*_min_, *p*_*l*_ · (*v*_*max*_ − *v*_min_)]. It is important to note that with *p*_*h*_ and *p*_*l*_ we redefine the ε parameter of the original iMAT formulation. The *ε* parameter of iMAT represented the minimum flux value attributed to highly expressed reactions (in iMAT: *ε* = 1 or any other user-defined flux value).
6. We maximize the number of lowly- (*r*_*L*_) and highly- (*r*_*H*_) expressed reactions whose flux levels are consistent with gene expression and can operate within the flux ranges defined in point 5. This optimization problem (OP) involves a mixed integer linear programming (MILP) formulation (see OP 1). We account for mass balances around every metabolite in the system at quasi-steady-state (eq. 1), where *S* is the stoichiometric matrix and *v* is the vector of reaction fluxes. We define the maximum (eq. 2) and minimum (eq. 3) flux requirement for the lowly- (*r*_*L*_) and highly- (*r*_*H*_) expressed reactions, respectively, as defined in point 5. In equation (2), the Boolean variable *y*_*i*_ takes the value 1 if reaction *i* is highly-expressed and forces the flux to be constrained between [*p*_*h*_ · (*v*_*max*_ − *v*_min_), *v*_*max*_]. The integer *y*_*i*_,.takes the value 1 (eq. 3) if reaction *i* is lowly-expressed and forces the flux to be constrained between [*v*_min_, *p*_*l*_ · (*v*_*max*_ − *v*_min_)]. The TEX-FBA can additionally account for thermodynamic constraints and metabolite concentrations (x_j_) as defined in TFA [25, 26] (eq. 5-7). We note that from the TEX-FBA formulation one can retrieve the original iMAT by setting the following parameters to *p*_*l*_ = 0 and *p*_*h*_ = 0.001 and the bounds to *v*_*min*,_ = 0 and *v*_*max*_ = 1000 (i.e. the original GEM flux upper bounds). Optimization objective:

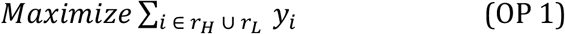

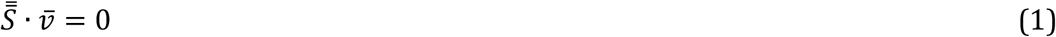

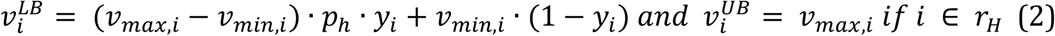

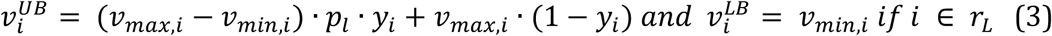

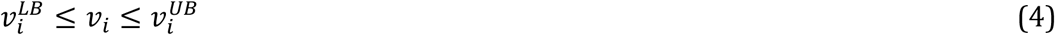

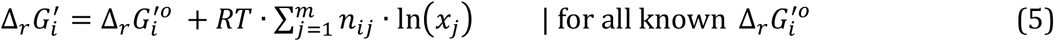

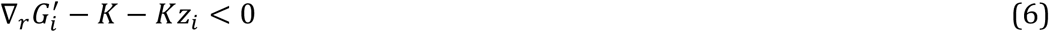

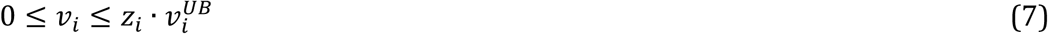

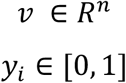
7. We define the solution of the TEX-FBA problem (OP 1) as a Maximum Consistency Score (MCS) between the reaction fluxes and gene expression. The MCS represents the maximum number of highly- and lowly-expressed reactions that can operate within the flux ranges defined in point 5. The highest possible value of the MCS is hence the total number of highly- (*r*_*H*_) and lowly- (*r*_*L*_) expressed reactions that we define as the Maximum Theoretical Consistency Score (MTCS) (Figure 5).

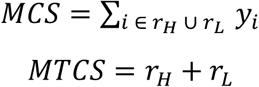
8. We integrate cut constraints [52] to identify all alternative flux profiles that result in the same MCS and hence are equally consistent with the experimental data.

**Figure 5.**
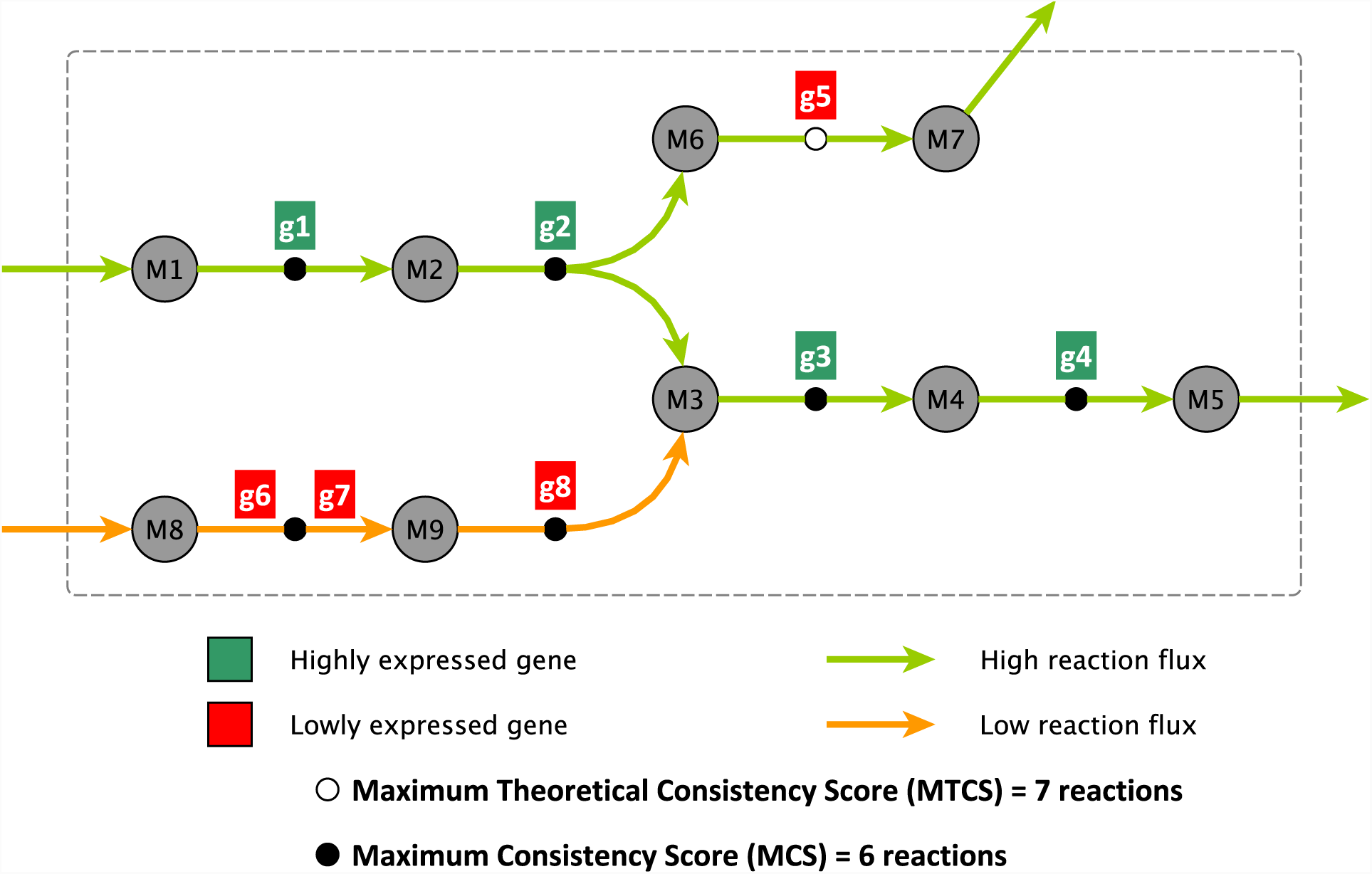
The TEX-FBA formulation (OP 1) applied to a toy model and definition of the Maximum Theoretical Consistency Score (MTCS) and Maximum Consistency Score (MCS). Metabolites (M) are shown with circles, reactions with arrows, and genes (g) with squares. Green is highly-expressed and red is lowly-expressed. In this figure, there are seven reactions with associated gene expression data, and the seven reactions are classified as highly- or lowly-expressed. The MTCS is hence seven. TEX-FBA (OP 1) indicates that all reactions but one (i.e., the one associated to g5) can operate in a flux range consistent with the gene expression. The MCS of the toy model with the give gene expression is thus six.

### Generation of alternative models at maximum consistency score

The presence of alternative solutions with the same MCS motivated us to study all alternative flux profiles at MCS. We generated one model per alternative solution with MCS. For this purpose, we set the binary variables (*y*_*i*_) of the OP 1 to the values that they obtained in each alternative solution. Thus, we applied flux constraints to the reactions with active binary variables (*y*_*i*_=1), as defined in the TEX-FBA OP 1.

### Flux Range Comparison (FRC) metric

The availability of a reference fluxomics dataset [38] motivated us to evaluate the performance of the models by comparing the predicted flux ranges (FRs) with the reference data. We computed a flux range comparison (FRC) metric to quantify the overlap and separation between the FRs of the 34 reactions for which we have C13 measurements [38] with the FRs obtained with FVA [40]:

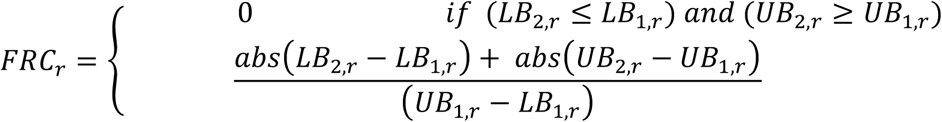

where *LB*_*1,r*_ and *LB*_*2,r*_ (and *UB*_*1,r*_ and *UB*_*2,r*_) are the lower (and upper) bound of flux ranges of a reaction *r* in the reference condition one and a condition two, respectively. In our study case, condition one involved the C13-based FRs and condition two represented the FVA-based FRs. The FRC of a reaction is zero when its FR is entirely inside the C13-based FR. The FRC is non-zero when a reaction FR has slight overlap or no overlap with the C13 FR. A high FRC value indicates poor agreement with C13-based flux measurement.

### Relative flexibility metric

We defined a relative flexibility metric to quantify the difference in flux ranges between two models, as done before [53, 54]. We determined the flux range of a reaction *i* in condition one 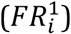 and condition two 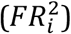 with a flux variability analysis (FVA) [40]:

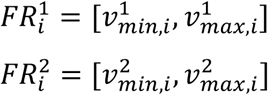

We then calculated the relative flexibility between the two conditions as follows:

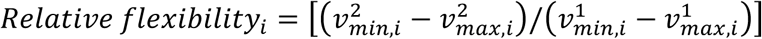

We computed the relative flexibility per reaction *i* for all reactions that carried flux in the reference FBA model. We also calculated an average relative flexibility (ARF) per metabolic subsystem and model for the XFBA, TFA, and XTFA models using the FBA model as a reference.

### Metabolic subsystem enrichment analysis

We performed a subsystem enrichment analysis to translate the information from TEX-FBA solutions at the reaction levels to the metabolic subsystems. We computed the hypergeometric probability density function (*P*) to quantify the probability of finding a metabolic subsystem with highly and lowly expressed reactions in a TEX-FBA solution:

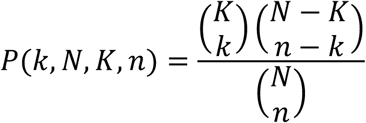

where *N* is the total number of reactions in the genome-scale model and *K* is the total number of reactions in a metabolic subsystem. The value of *n* indicates the number of reactions with gene expression constraints in a solution from TEX-FBA (OP 1), and *k* is the number of reactions with gene expression constraints in a solution from TEX-FBA (OP 1) that belong to the subsystem of interest.

### Reaction and gene essentiality for growth

We performed reaction and gene knockouts *in silico* following the Fast-SL approach [55] with FBA [22] and TFA [25, 26]. We considered a reaction or gene is essential for growth when the growth rate upon its knockout is below 10% of the wild-type growth rate. We applied the same approach to all the models used in this study to identify the essential genes and reactions for growth.

### Software usage and settings

We solved FBA [22] with the version of the COBRA Toolbox v2.0 [56] included in the matTFA toolbox [50]. We performed TFA [25, 26] with the matTFA toolbox [50]. We used MATLAB (R2016a and R2017a) and CPLEX (ILOG IBM v12.51 and v12.71) as the linear solver.

### Software availability

Documented implementation of TEX-FBA framework in MATLAB is available on www.github.com/EPFL-LCSB/. The software package includes a tutorial that indicates step-by-step how to integrate gene expression information into a genome-scale model using TEX-FBA. The genome-scale model of *E. coli* iJO1366 [39] and data [38] used in this study are also available. The tutorials indicate how to reproduce the results and figures for this paper. TEX-FBA requires the matTFA toolbox [50] for TFA.

Author Note
VP, DHG, ACP, and VH are supported by the École Polytechnique Fédérale de Lausanne (EPFL). VP, ACP, and VH are supported by the RTD grant MalarX within SystemsX.ch, the Swiss Initiative for Systems Biology evaluated by the Swiss National Science Foundation.

